## supplementary tables and figures for "IgG1 responses following SARS-CoV-2 infection are polyclonal and highly personalized, whereby each donor and each clone displays a distinct pattern of cross-reactivity against SARS-CoV-2 variants"

**Supplementary Table 1| Plasma donor characteristics**.

| Record Id | Proven PCR test of infection | Variant of infection | Age | Gender | Severity score WHO | Respiratory symptoms 1=Yes; 2=No | Fever 1=Yes; 2=No | Hospital admission 1=Yes; 2=No | IC 1=Yes; 2=No | Duration admission in days | Chronic immune suppresion 1=Yes; 2=No | Short immune suppresion during COVID-19? | Treatment during hospital admission? | Date when COVID-19 symptoms started | Date of blood withdrawal | Days between symptoms and blood withdrawal |
|---|---|---|---|---|---|---|---|---|---|---|---|---|---|---|---|---|
| **002** | Yes | WT | 44 | Female | 2 | 1 | 1 | 2 | 2 | 0 | No | No | | 2020-02-25 | 2020-03-23 | 27 |
| **003** | Yes | WT | 69 | Male | 4 | 1 | 1 | 1 | 1 | 13 | No | No | Cefotaxim, Ciproflox | 2020-03-08 | 2020-03-23 | 15 |
| **303** | Yes | α | 66 | Male | 2 | 1 | 1 | 2 | 2 | 0 | No | No | | 2021-01-08 | 2020-02-04 | 27 |
| **304** | Yes | α | 59 | Female | 2 | 1 | 1 | 2 | 2 | 0 | No | No | | 2021-01-02 | 2020-02-04 | 33 |
| **307** | Yes | β | 18 | Female | 1 | 1 | 1 | 2 | 2 | 0 | No | No | | 2021-01-13 | 2021-02-12 | 30 |
| **308** | Yes | β | 18 | Male | 1 | 1 | 1 | 2 | 2 | 0 | No | No | | 2021-01-07 | 2021-02-12 | 36 |
| **309** | Yes | γ | 32 | Female | 1 | 1 | 2 | 2 | 2 | 0 | No | No | | 2021-01-09 | 2021-02-19 | 41 |
| **310** | Yes | γ | 37 | Male | 1 | 1 | 2 | 2 | 2 | 0 | No | No | | 2021-01-12 | 2021-02-19 | 38 |

**Supplementary Table 2| The total amount of S-protein directed IgG1 clones, and their concentrations vary substantially in between donors.** *For each donor and each variant, the number of IgG1 clones detected binding to each VOC S-protein-trimer mutant is provided and compared to the total number of IgG1 clones detected in full plasma (bottom rows). Making use of the two recombinant IgG1 mAbs internal standards, we could also estimate the total and individual concentrations of IgG1s. As shown in the table the total concentration of IgG1s in these eight donors varies in between 100 ug/ml to 582 ug/ml. The total concentrations of IgG1s binding to the VOC S-protein-trimer mutants, varies much more from 0.1 ug/ml to 31,6 mg/mL.*

| | | 002 | 003 | 303 | 304 | 307 | 308 | 309 | 310 |
|---|---|---|---|---|---|---|---|---|---|
| | VOC | WT | WT | α | α | β | β | γ | γ |
| | Score | 2 | 4 | 2 | 2 | 1 | 1 | 1 | 1 |
| WT | # of clones | 46 | 192 | 277 | 13 | 2 | 5 | 71 | 23 |
| | Concentration (µg/mL) | 0.6 | 21.7 | 31.6 | 0.6 | <0.1 | 0.3 | 2.3 | 0.3 |
| α | # of clones | 9 | 116 | 281 | 19 | 1 | 4 | 50 | 27 |
| | Concentration (µg/mL) | 0.1 | 12.3 | 14.6 | 0.7 | <0.1 | 0.1 | 0.8 | 0.3 |
| β | # of clones | 9 | 68 | 204 | 6 | 3 | 8 | 39 | 18 |
| | Concentration (µg/mL) | 0.1 | 7.0 | 10.1 | 0.6 | <0.1 | 0.3 | 1.2 | 0.3 |
| γ | # of clones | 7 | 114 | 208 | 11 | 3 | 5 | 82 | 33 |
| | Concentration (µg/mL) | 0.2 | 6.7 | 9.5 | 0.9 | <0.1 | 0.3 | 3.1 | 0.5 |
| | # of clones | 381 | 370 | 517 | 247 | 363 | 386 | 454 | 284 |
| | Concentration (µg/mL) | 356.9 | 582.1 | 317.8 | 100.3 | 219.1 | 272.8 | 340.7 | 153.7 |

**A**

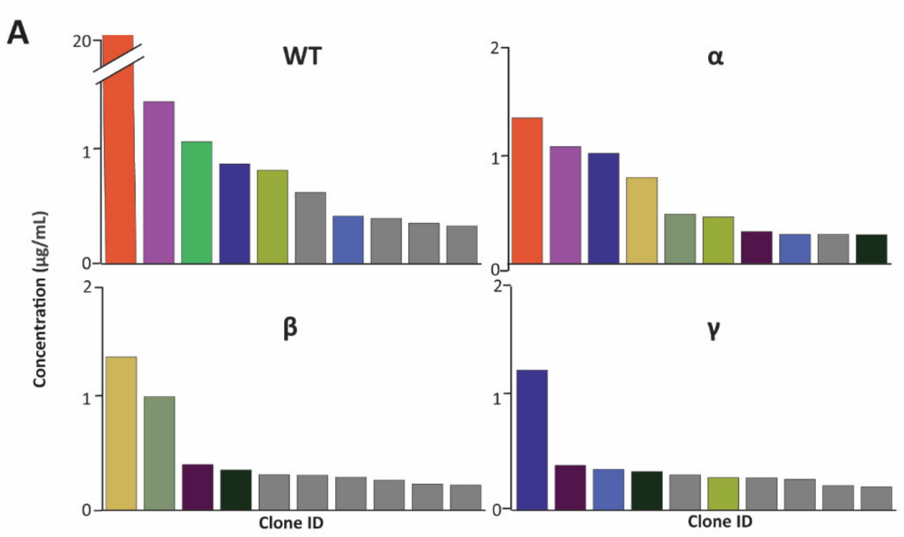


**B**

| Clone ID (RT (min) # Mass (Da)) | WT (µg/mL) | α (µg/mL) | β (µg/mL) | γ (µg/mL) |
|---|---|---|---|---|
| 20.1 **449** 48,655.2 | 18.1 | 1.3 | 0.2 | 0.1 |
| 18.4 **208** 47,361.0 | 1.4 | 1.0 | N.D. | N.D. |
| 18.0 **124** 46,780.0 | 0.9 | 1.0 | N.D. | 1.2 |
| 24.9 **17** 50,677.2 | 0.1 | 0.8 | 1.3 | 0.1 |
| 27.2 **442** 48,315.6 | 0.1 | 0.4 | 1.0 | N.D. |
| 15.0 **354** 47,814.5 | 0.8 | 0.4 | N.D. | 0.3 |
| 22.2 **387** 49,627.9 | 0.1 | 0.3 | 0.4 | 0.4 |
| 18.0 **483** 48,880.5 | 0.4 | 0.3 | 0.2 | 0.4 |
| 16.3 **456** 48,618.2 | 0.1 | 0.3 | 0.4 | 0.3 |
| 21.6 **455** 48,623.5 | 1.1 | 0.1 | N.D. | 0.3 |

**C**

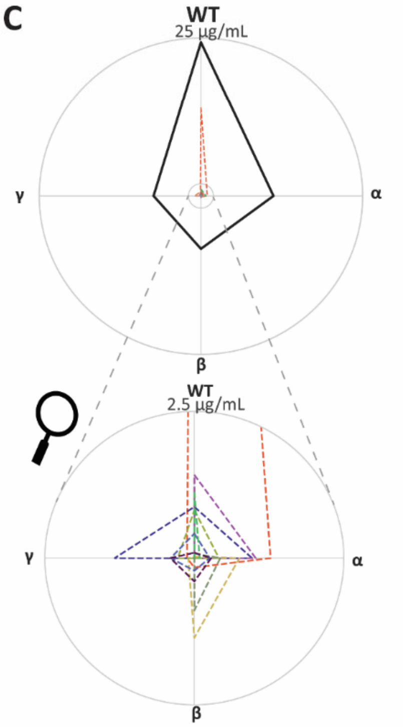


**Supplementary Figure 1| Within a single donor (303) IgG1 clones display distinctive cross-reactivity versus the S-protein variants. A.** *Quantitative comparison of the ten most abundant IgG1 fab clones enriched from the plasmaof donor 003. Each bar represents one of the 10 most abundant antibody clones with the height indicating concentration in ug/ml. using the same colors for the clones corresponding to panel B. Each clone that was not in the total top 10 but was in the top 10 for that specific VOC is colored grey* **B.** *The table depicts the top 10 most abundant clones showing the concentration that in total is enriched using the different S-protein variants for that specific clone. This top 10 was selected based on the clones that in total, by summing up the concentrations found against the different S-protein variants, showed the highest concentrations. The colored bars in (A) and the dotted lines in the radar plot in (C) are corresponding to the clones in the table.* **C.** *Radar plot with on each edge one of the tested S-protein variants. These plots depict the difference in binding of specific clones against the different S-protein variants. The thick solid gray line representing the sum of all enriched IgG1 clones against the specific S-protein variants. The color coding is identical as used in (A) and (B), corresponding to the same unique IgG1 clones.*

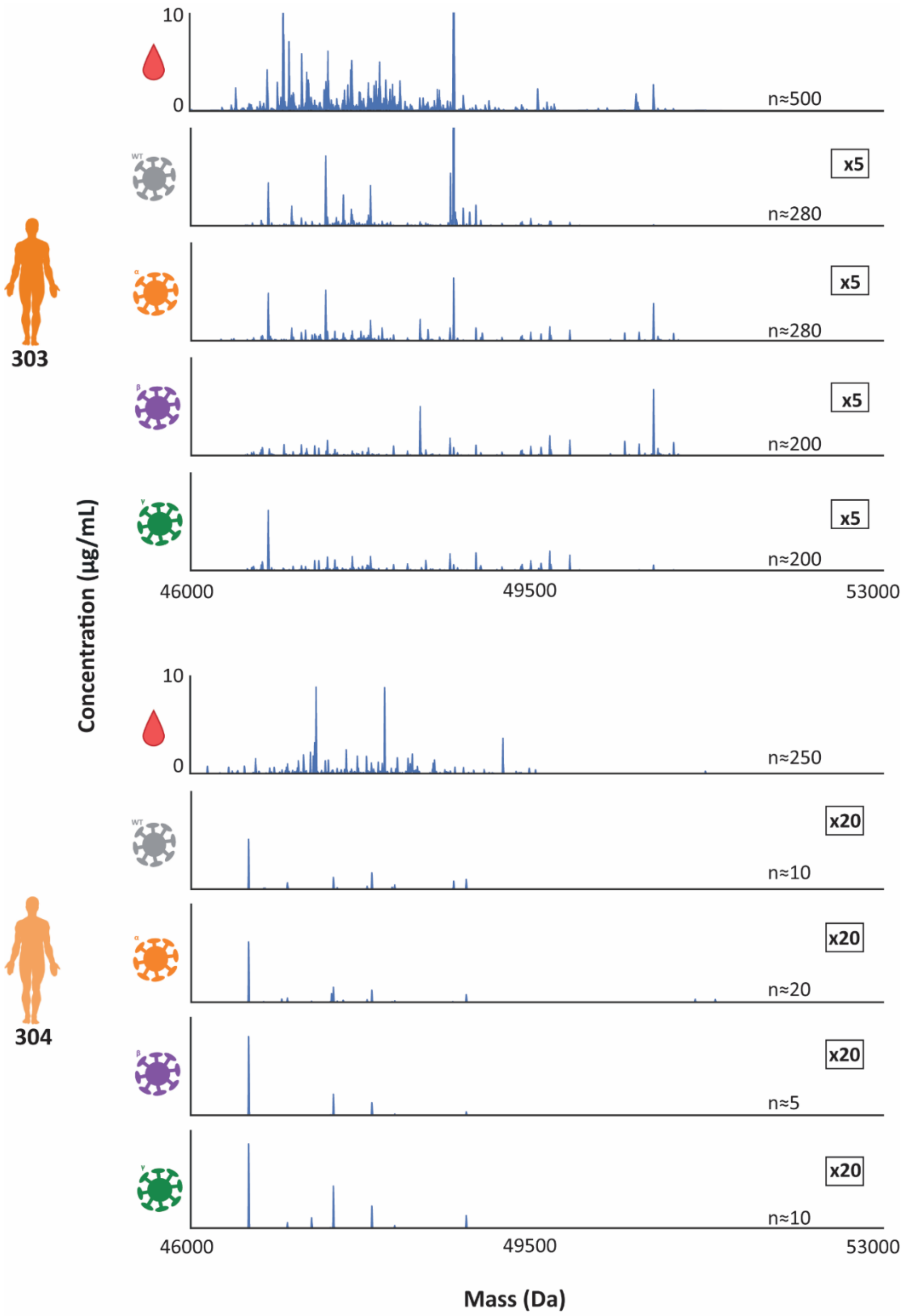


**Supplementary Figure 2| Fab mass profiles for donor 303 and 304.** *From top to bottom, the full plasma Fab profile, followed by the S-protein directed Fab profiles using the WT, Alpha, Beta and Gamma VOC variant. Each peak represents a unique Fab at its detected mass and plasma concentration. In each plot the number of unique Fab identified is indicated. The number on the right of each S-protein directed Fab profile, shows the magnification of the y-axis, compared to the data for the full plasma Fab profile.*

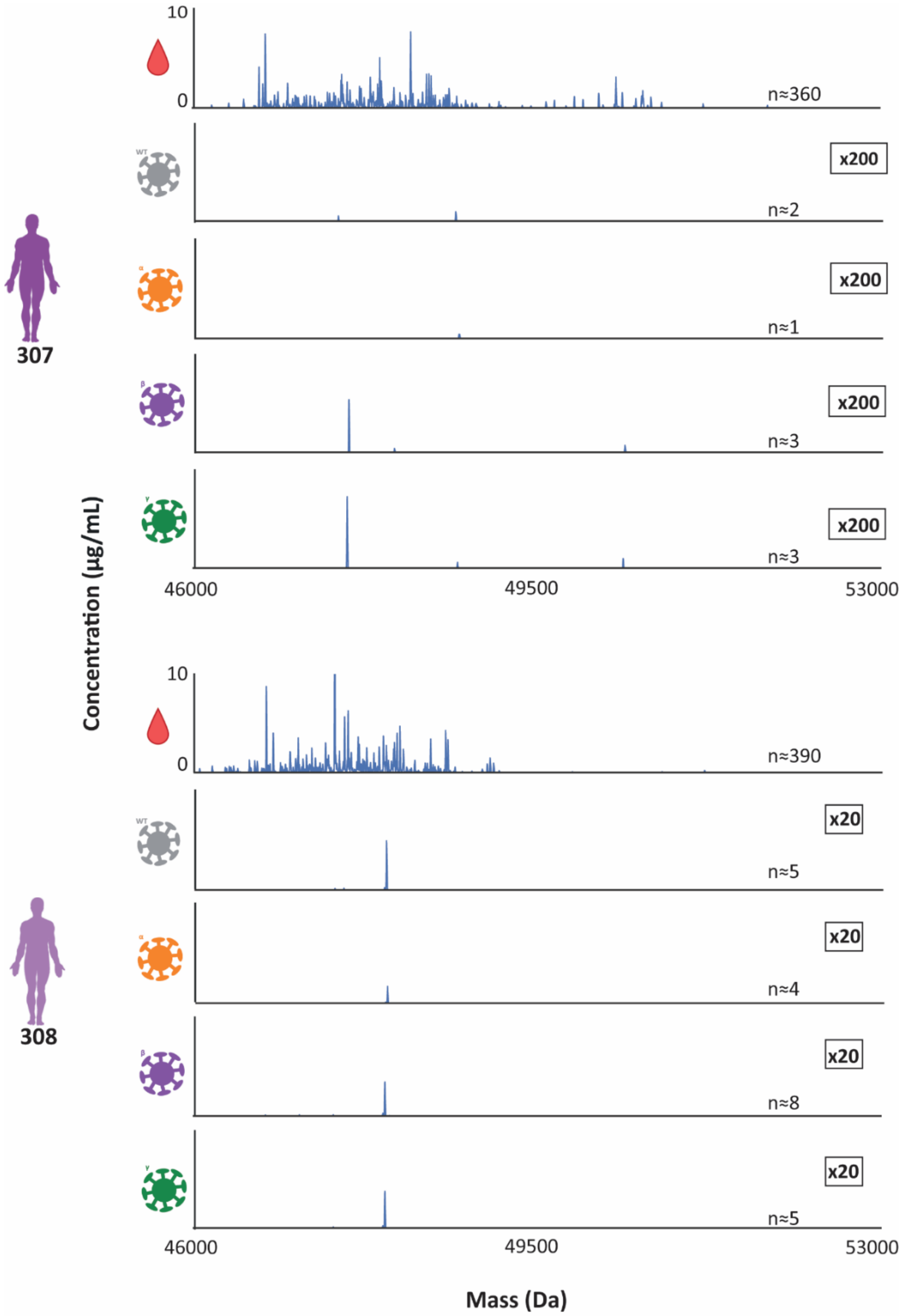


**Supplementary Figure 3| Fab mass profiles for donor 307 and 308.** *From top to bottom, the full plasma Fab profile, followed by the S-protein directed Fab profiles using the WT, Alpha, Beta and Gamma VOC variant. Each peak represents a unique Fab at its detected mass and plasma concentration. In each plot the number of unique Fab identified is indicated. The number on the right of each S-protein directed Fab profile, shows the magnification of the y-axis, compared to the data for the full plasma Fab profile.*

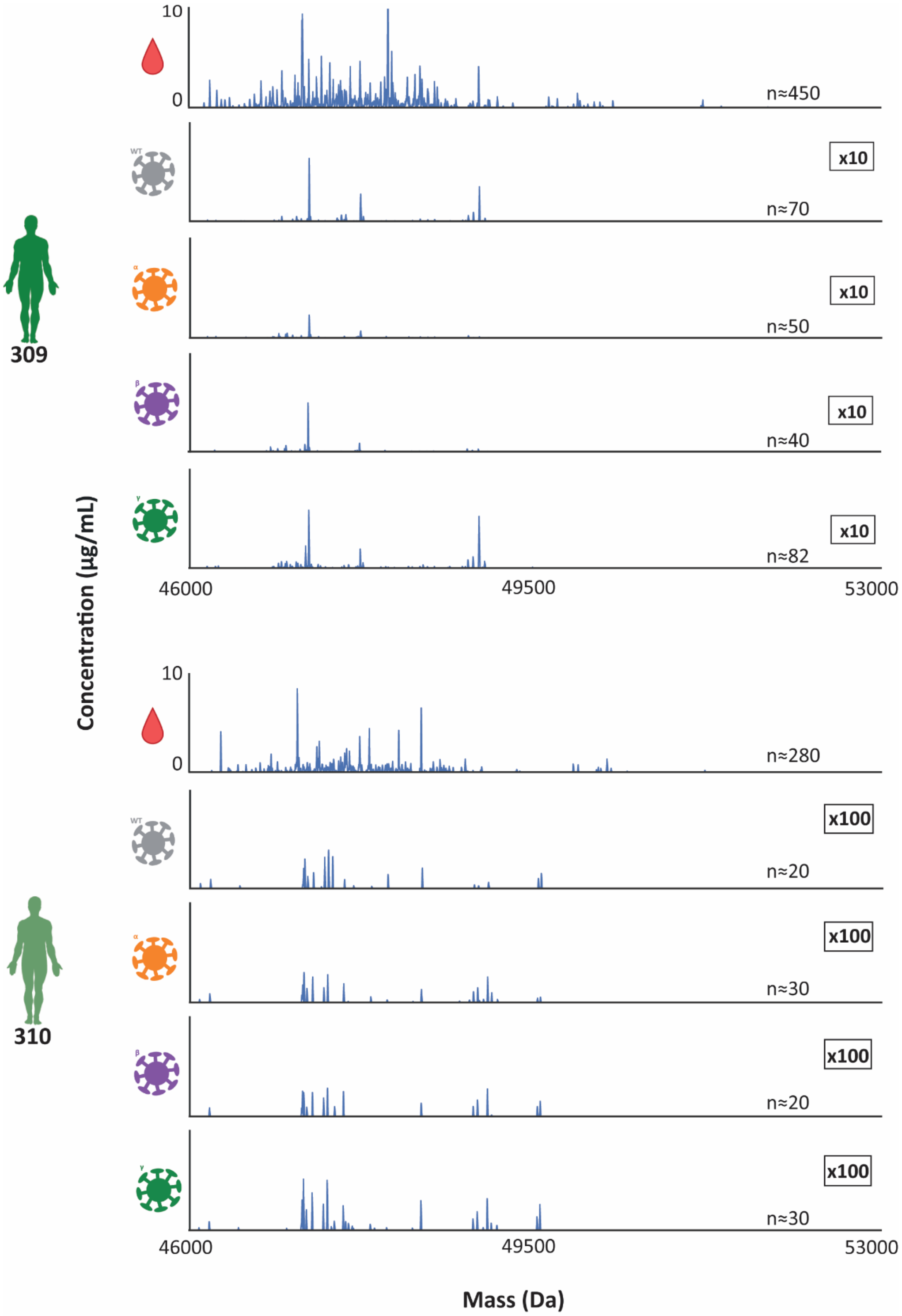


**Supplementary Figure 4| Fab mass profiles for donor 309 and 310.** *From top to bottom, the full plasma Fab profile, followed by the S-protein directed Fab profiles using the WT, Alpha, Beta and Gamma VOC variant. Each peak represents a unique Fab at its detected mass and plasma concentration. In each plot the number of unique Fab identified is indicated. The number on the right of each S-protein directed Fab profile, shows the magnification of the y-axis, compared to the data for the full plasma Fab profile.*
